# BST-2 inhibits SARS-CoV-2 egress at intracellular membranes and is neutralized by ORF7a

**DOI:** 10.1101/2025.07.07.663519

**Authors:** Adam Smith, Xinhong Dong

## Abstract

Bone marrow stromal antigen 2 (BST-2, or tetherin) is an interferon-inducible host restriction factor that inhibits the release of enveloped viruses by tethering nascent virions to cellular membranes. While its antiviral function is well established in retroviral systems, its role in SARS-CoV-2 egress remains unclear. Here, we used a virus-like particle (VLP) system composed of SARS-CoV-2 structural proteins M, E, and N to investigate the impact of BST-2 on viral particle release. BST-2 significantly inhibited VLP release in HEK293T and Calu-3 lung epithelial cells. Confocal microscopy revealed that BST-2 colocalizes with viral structural proteins at the endoplasmic reticulum-Golgi intermediate compartment (ERGIC), the main site of coronavirus assembly. We next evaluated the roles of the SARS-CoV-2 accessory proteins ORF3a and ORF7a in overcoming this restriction. ORF3a localized to endolysosomal compartments and promoted VLP release through a BST-2-independent mechanism, without altering BST-2 expression or localization. In contrast, ORF7a colocalized with both BST-2 and ERGIC markers and restored VLP release by promoting BST-2 degradation. Notably, ORF7a also relieved BST-2-mediated restriction of HIV-1 VLP release, suggesting a conserved antagonistic function. These findings identify BST-2 as an intracellular inhibitor of SARS-CoV-2 particle release and establish ORF7a as viral accessory antagonist that neutralizes this host defense.

## Introduction

Bone marrow stromal antigen 2 (BST-2), also known as tetherin or CD317, is an interferon-inducible transmembrane protein that acts as a broad-spectrum antiviral restriction factor. It inhibits the release of a wide range of enveloped viruses by physically tethering budding virions to the host cell membrane, thereby limiting viral dissemination and replication[1]. BST-2 is structurally unique among restriction factors, comprising an N-terminal cytoplasmic tail, a single transmembrane domain, an extended extracellular coiled-coil domain that mediates dimerization, and a C-terminal glycosylphosphatidylinositol (GPI) anchor. This dual membrane-anchoring configuration allows BST-2 to bridge host and viral membranes, with one end inserted into cell membranes and the other into the viral envelope, effectively trapping virions at the cell surface or within intracellular compartments[2, 3].

BST-2 is constitutively expressed in various cell types and is strongly upregulated by type I and type III interferons. In addition to its presence at the plasma membrane, BST-2 localizes to intracellular compartments including endosomes, the trans-Golgi network (TGN), and the endoplasmic reticulum (ER), consistent with its role in restricting viruses that assemble at different subcellular sites[4]. Initially characterized as a restriction factor for HIV-1, BST-2 has since been shown to inhibit a wide range of viruses, including Ebola, Marburg, Lassa, influenza, Nipah, herpesviruses, and others [3, 5–9].

Recent studies suggest that BST-2 also restricts coronaviruses. BST-2 has been reported to inhibit SARS-CoV-1 particle release [10], sequester human coronavirus 229E (HCoV-229E) in intracellular vesicles [11], and mediate lysosomal degradation of nucleocapsid protein in porcine endemic diarrhea virus (PDEV) [12]. These findings imply that BST-2 may act at multiple stages of the coronavirus life cycle. However, its role in restricting SARS-CoV-2, the causative agent of COVID-19, remains incompletely defined.

SARS-CoV-2 is a positive-sense, single-stranded RNA virus that belongs to the *Betacoronavirus* genus. In contrast to many enveloped viruses that bud at the plasma membrane, SARS-CoV-2 assembles and buds at ER-Golgi intermediate compartment (ERGIC), a specialized organelle located between the rough ER and the cis-Golgi [13–17]. Structural proteins including membrane (M), envelope (E), nucleocapsid (N), and spike (S) are synthesized and trafficked to the ERGIC, where virion assembly occurs. The M protein plays a central role in coordinating assembly, interacting with E and S in the membrane and recruiting the N-RNA complex. E protein facilitates membrane curvature and scission, while S mediates receptor binding and entry. Fully assembled virions are transported through the Golgi and released by exocytosis [16,17].

Although this intracellular route of assembly and egress is well established, it is not yet known whether BST-2 restricts SARS-CoV-2 at these intracellular sites, or whether the virus has evolved mechanisms to antagonize this restriction [18, 19].

Several viruses have developed specific proteins that counteract BST-2. For example, the HIV-1 accessory protein Vpu induces BST-2 degradation through β-TrCP-mediated ubiquitination and lysosomal targeting [20, 21], and also impairs BST-2 trafficking to viral budding sites [22, 23]. Similar antagonistic strategies are used by other viruses such as Ebola virus glycoprotein [24] and K5 ubiquitin ligase of Kaposi’s sarcoma-associated herpesvirus (KSHV) [25]. In coronaviruses, SARS-CoV-1 spike (S) protein and ORF7a have been shown to modulate BST-2 function by promoting degradation or interfering with glycosylation [10]. In SARS-CoV-2, both the S protein and ORF7a have been proposed as potential BST-2 antagonists, but their exact roles remain poorly characterized [26]. Additionally, the SARS-Cov-2 accessary protein ORF3a has been implicated in modulating membrane trafficking and autophagy, raising the possibility that it could indirectly influence BST-2 activity [27].

In this study, we used a SARS-Cov-2 virus-like particle (VLP) system composed of structural proteins M, E, and N to investigate whether BST-2 restricts viral egress and whether SARS-CoV-2 accessory proteins counteract this restriction. We show that BST-2 colocalizes with structural proteins at the ERGIC and significantly inhibits VLP release in both HEK 293T and Calu-3 lung epithelial cells. Furthermore, we demonstrate that ORF3a enhances VLP release through a mechanism that is independent of BST-2, whereas ORF7a colocalizes with BST-2 and restores VLP release by promoting its degradation. These findings identify the ERGIC as a novel site of BST-2-mediated restriction and establish ORF7a as a viral antagonist that facilitates SARS-CoV-2 egress.

## Results

### SARS-CoV-2 structural proteins accumulate at the ERGIC compartment

To define the subcellular localization of SARS-CoV-2 structural proteins and assess their targeting to the ERGIC, the primary site of coronavirus assembly and budding, we performed high-resolution confocal microscopy in HeLa cells transiently transfected with plasmids encoding individual viral structural proteins. Enhanced green fluorescent protein (EGFP)-tagged versions of the E and M proteins were used to enable direct fluorescence imaging, while the N and S proteins were expressed without tags and detected via immunofluorescence using specific antibodies. At 48 hours post-transfection, cells were fixed and permeabilized, followed by staining with anti-ERGIC-53, a canonical marker of the ERGIC that localizes to perinuclear membranes and is commonly used to define this compartment. For detection of untagged proteins, anti-N and anti-S antibodies were used in conjunction with appropriate fluorophore-conjugated second antibodies. All four structural proteins displayed a predominantly perinuclear distribution consistent with the known localization of the ERGIC. EGFP-E and EGFP-M exhibited a reticular and punctate pattern tightly colocalized with ERGIC-53-positive membranes. Similarly, both N and S proteins showed strong perinuclear enrichment and colocalized with ERGIC-53 as determined by immunofluorescence. Quantitative analysis using Pearson’s correlation coefficient confirmed a high degree of colocalization between each structural protein and ERGIC-53: EGFP-E (R = 0.78 ± 0.03), EGFP-M (R = 0.75 ± 0.06), N (R = 0.64 ± 0.08), and S (R = 0.65 ± 0.13) (Fig. 1a). These data indicate that SARS-CoV-2 structural proteins are efficiently targeted to the ERGIC in HeLa cells and support the conserved role of this compartment as the intracellular site for coronavirus virion assembly.

**Figure 1.**
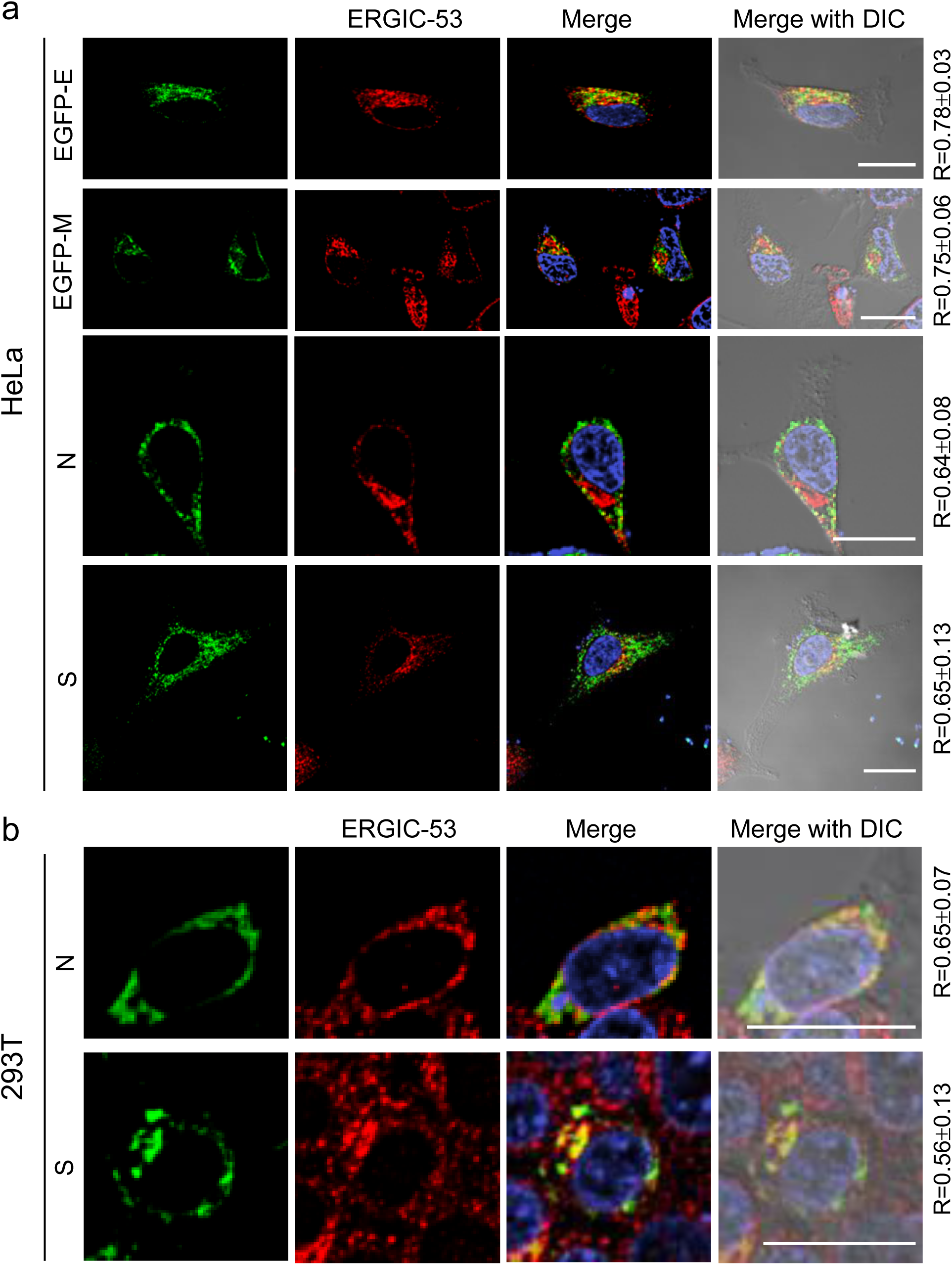
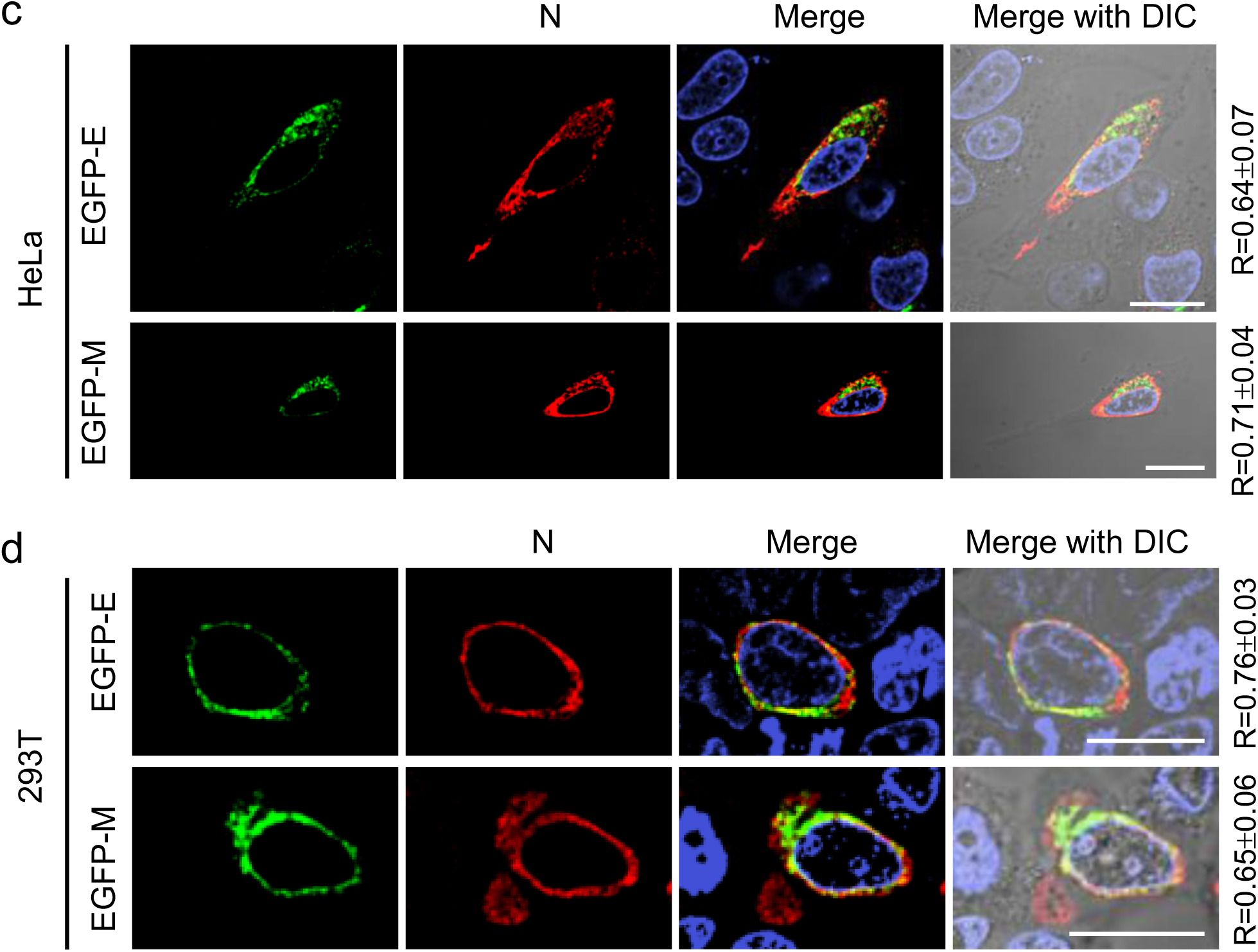
SARS-CoV-2 structural proteins localize to the ERGIC and colocalize with each other. **(a)** HeLa cells were transfected with plasmids encoding EGFP-E (top row), EGFP-M (second row), N (third row), or S (bottom row). At 48 h post-transfection, cells were fixed, permeabilized, and stained with anti-ERGIC-53 alone (top two rows), anti-N and anti-ERGIC-53 (third row), or anti-S and anti-ERGIC-53 (bottom row). EGFP-E, EGFP-M, N, or S proteins are shown in green (far-left panels); EGRIC-53 is shown in red (left panels). Merged images with DAPI nuclear staining are shown in the right panels, and merged DAPI/DIC images in the far-right panels. **(b)** HEK 293T cells were transfected with plasmids encoding N (top row) or S (bottom row) and stained with anti-ERGIC-53 and either anti-N or anti-S. Viral proteins are shown in green; ERGIC-53 in red. Merged images with DAPI and DAPI/DIC are shown in the right and far-right panels. **(c, d)** HeLa (c) or HEK 293T (d) cells were co-transfected with EGFP-E and N (top rows) or EGFP-M and N (bottom rows), followed by anti-N staining. EGFP-tagged proteins are shown in green; N is shown in red. Merged DAPI and DAPI/DIC images are shown in the right and far-right panels. Scale bars, 20 μm. Colocalization was quantified using Pearson’s correlation coefficient (R). Data represent means ± SD from 20-30 cells per condition, based on ≥ 3 independent experiments.

To determine whether the subcellular targeting of structural proteins to the ERGIC is modulated by BST-2, we repeated the localization analysis in HEK 293T cells, which lack endogenous BST-2 expression. In these cells, both N and S proteins retained substantial colocalization with ERGIC-53, with Pearson’s R values of 0.65 ± 0.07 and 0.56 ± 0.13, respectively (Fig. 1b). This finding suggests that ERGIC localization of SARS-CoV-2 structural proteins occur independently of BST-2 expression and is conserved feature across distinct human cell lines.

To further assess the potential for coordinated trafficking or complex formation among structural proteins during assembly, we examined colocalization between EGFP-E or EGFP-M and the N protein. In HeLa cells co-expressing EGFP-E or EGFP-M with untagged N proteins, we observed robust colocalization in perinuclear regions. Quantitative analysis yielded Person’s correlation coefficients of 0.64 ± 0.07 for EGFP-E/N and 0.71 ± 0.04 for EGFP-M/N (Fig. 1c). A similar pattern was observed in HEK 293T cells, where EGFP-E and EGFP-M colocalized with N at R values of 0.76 ± 0.03 and 0.65 ± 0.06, respectively (Fig. 1d). These data are consistent with the hypothesis that structural proteins not only converge at the ERGIC but may also interact directly or indirectly to facilitate assembly of virion components into budding particles.

Collectively, these results establish that the SARS-CoV-2 E, M, N, and S structural proteins accumulate at the ERGIC regardless of cell type or BST-2 expression status. The observed spatial convergence of these proteins supports the ERGIC as the central platform for coronavirus assembly and highlights the potential for coordinated trafficking and protein-protein interactions as critical steps in virion morphogenesis.

### BST-2 restricts the release of SARS-CoV-2 virus-like particle release at the ERGIC

We next asked whether BST-2 impairs the release of SARS-CoV-2 virus-like particles (VLPs). To investigate this, we employed a VLP production system in HEK 293T cells, in which co-expression of the SARS-CoV-2 E, M, and N proteins is sufficient to drive the assembly and secretion of VLPs in the absence of genomic RNA or accessory proteins [28–30]. Consistent with previous studies, the co-expression of E, M, and N proteins in HEK 293T cells led to efficient production and release of VLPs into the culture supernatant. Western blot analysis of VLP-containing supernatants confirmed the presence of viral structural proteins, indicating successful particle formation. To assess the impact of BST-2, we co-transfected cells with a plasmid encoding FLAG-tagged human BST-2. Remarkably, BST-2 expression resulted in a ∼3-fold reduction in VLP release compared to vector control, as quantified by densitometric analysis of Western blots (Fig. 2a). This significant inhibition suggests that BST-2 retains its antiviral activity against SARS-CoV-2 structural components, even in the absence of other viral or host factors.

**Figure 2.**
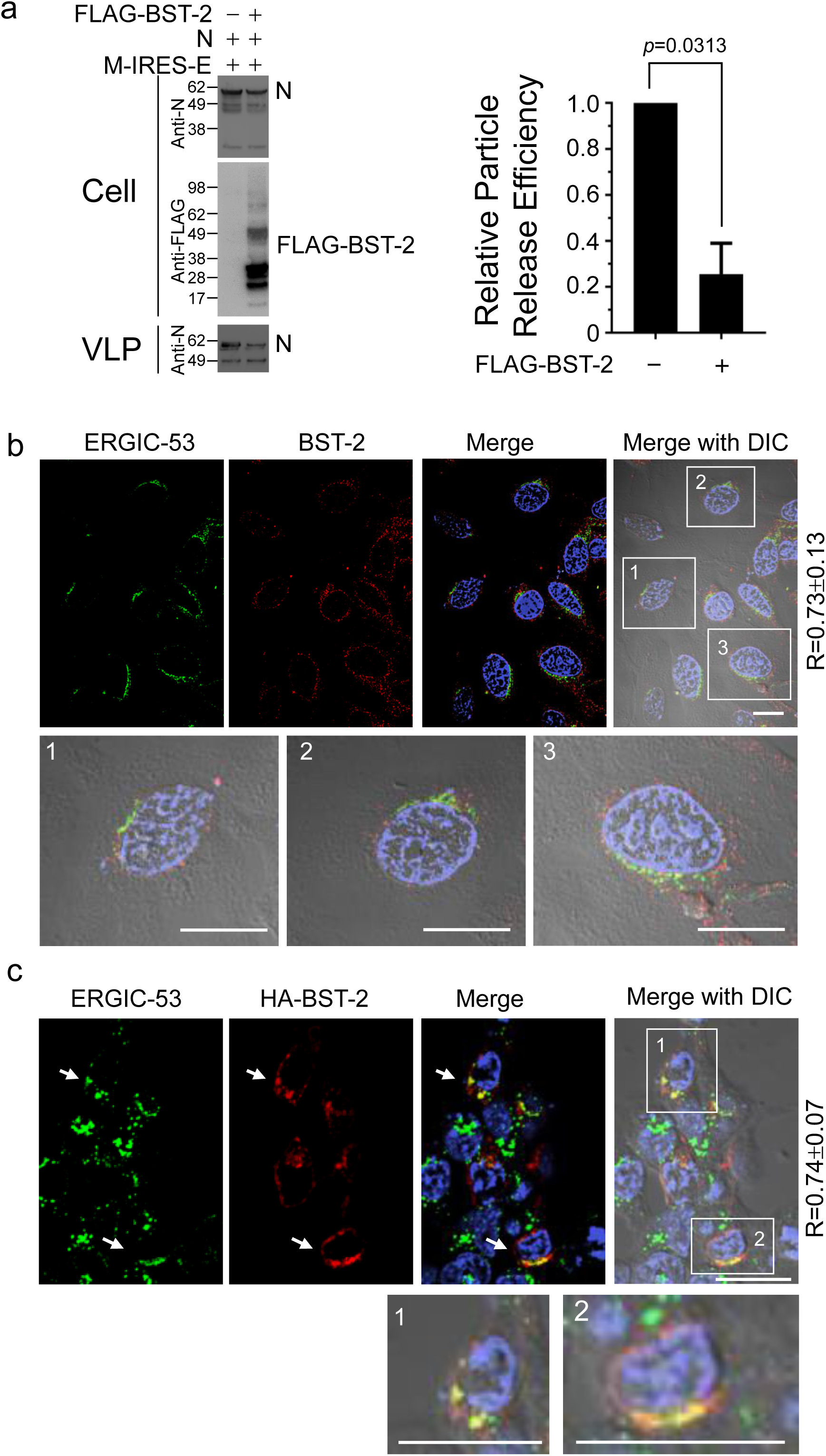
BST-2 restricts SARS-CoV-2 VLP release and colocalizes with the ERGIC. **(a)** HEK 293T cells were co-transfected with M-IRES-E, N, and either empty vector or FLAG-BST-2. At 48 h post-transfection, cell lysates and pelleted VLPs were analyzed by western blot using anti-FLAG and anti-N antibodies. Relative particle release efficiency was quantified by normalizing N protein levels in supernatants from FLAG-BST-2-expressing cells to vector control (set to 1.0). Right graph shows means ± SD from 3 independent experiments. **(b)** HeLa cells were stained with anti-ERGIC-53 and anti-BST-2. ERGIC-53 is shown in green; BST-2 in red. Merged images with DAPI and DAPI/DIC are shown in the right and far-right panels, respectively. Enlarged views of boxed regions are shown below. **(c)** HEK 293T cells expressing HA-BST-2 were staining with anti-HA and anti-ERGIC-53. ERGIC-53 is shown in green; HA-BST-2 in red. Merged DAPI/DIC images and enlarged insets are shown. Scale bars, 20 μm. Pearson’s R values represent means ± SD from ≥ 3 experiments, analyzing 20-30 cells per condition.

Although BST-2 is canonically localized to the plasma membrane, endosomal compartments, and the trans-Golgi network (TGN), its ability to inhibit SARS-CoV-2 release suggested it might also function at the site of virion assembly-the ERGIC. To assess this possibility, we performed confocal immunofluorescence microscopy in HeLa cells and probed for endogenous BST-2 and ERGIC-53. We observed pronounced perinuclear colocalization between BST-2 and ERGIC-53, with a Pearson’s correlation coefficient of R = 0.73 ± 0.13 (Fig. 2b), indicating a substantial proportion of endogenous BST-2 resides at the ERGIC.

To confirm these findings and assess localization in the context of overexpression, we performed a similar analysis in HEK 293T cells transfected with HA-tagged BST-2. In these cells, HA-BST-2 also showed robust perinuclear staining and colocalized strongly with ERGIC-53 (R = 0.74 ± 0.07) (Fig. 2c), supporting the idea that BST-2 can access the ERGIC under both endogenous and overexpression conditions. This previously underappreciated localization pattern provides a mechanistic basis for how BST-2 might restrict SARS-CoV-2 egress. By localizing to the ERGIC where E, M, and N coalesce to assemble VLPs, BST-2 is positioned to physically tether nascent particles and prevent their release into the secretory pathway.

Together, these findings identify BST-2 as a potent inhibitor of SARS-CoV-2 VLP release and reveal that a functionally relevant pool of BST-2 localizes to the ERGIC. This expands the known antiviral repertoire of BST-2 beyond its classical role at the plasma membrane and highlights a potential mechanism by which host cells can restrict coronavirus egress at intracellular assembly sites.

### BST-2 colocalizes with SARS-CoV-2 structural proteins at Intracellular and plasma membranes

To determine whether BST-2 physically associates with viral structural proteins, we performed quantitative colocalization analyses in HeLa cells transiently transfected with EGFP-tagged E or M, or untagged N. Immunostaining for endogenous BST-2 revealed that BST-2 localized predominantly to perinuclear and vesicular compartments, overlapping moderately with each of the structural proteins. Person’s correlation coefficients indicated moderate colocalization with EGFP-M (R = 0.57 ± 0.06), EGFP-E (R = 0.59 ± 0.08), and N (R = 0.54 ± 0.03) (Fig. 3a). These correlation values suggest that endogenous BST-2 and viral structural proteins are co-resident within common intracellular compartments, including the ERGIC and downstream trafficking intermediates.

**Figure 3.**
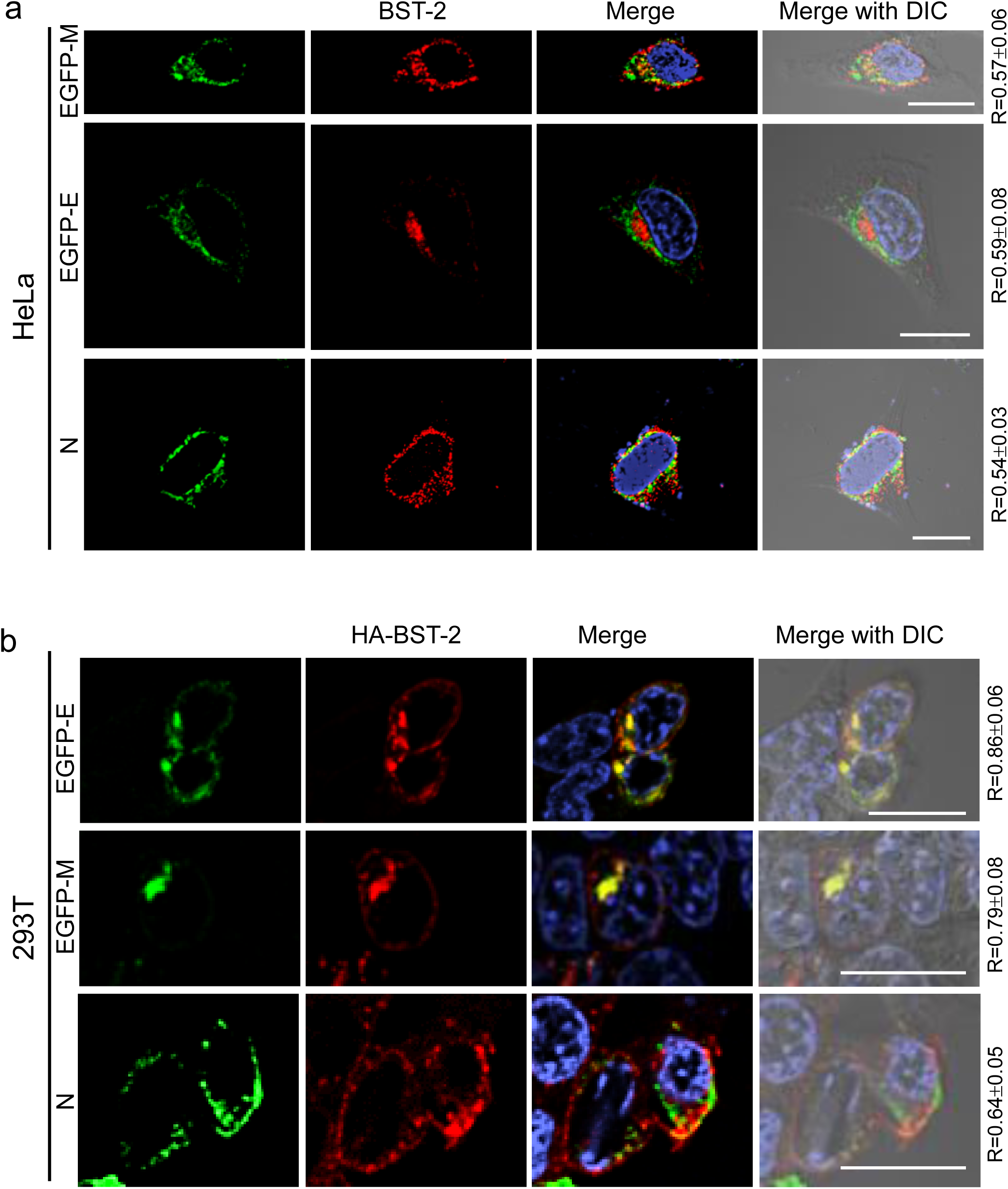
SARS-CoV-2 structural proteins colocalize with BST-2. **(a)** HeLa cells were transfected with EGFP-M (top), EGFP-E (middle), or N (bottom), and stained with anti-BST-2 (top two rows) or anti-BST-2 and anti-N (bottom row). Viral proteins are shown in green; BST-2 in red. Merged DAPI and DAPI/DIC images are shown in the right and far-right panels. **(b)** HEK 293T cells were co-transfected with HA-BST-2 and EGFP-E (top), EGFP-M (middle), or N (bottom), followed by staining with anti-HA (top two rows) or anti-HA and anti-N (bottom row). Viral proteins are shown in green; HA-BST-2 in red. Scale bars, 20 μm. Pearson’s R values represent means ± SD from ≥ 3 independent experiments.

To further assess the extent and subcellular context of these associations, we repeated the analysis in HEK 293T cells transiently transfected with HA-tagged human BST-2. As previously reported, exogenously expressed BST-2 localizes to both intracellular membranes and the plasma membrane. In our system, it exhibited stronger colocalization with viral structural proteins, consistent with its broader subcellular distribution and elevated expression levels. Quantitative analysis revealed strong colocalization of exogenous BST-2 with EGFP-E (R = 0.86 ± 0.06), EGFP-M (R = 0.79 ± 0.08), and N (R = 0.64 ± 0.05) (Fig. 3b). This enhanced colocalization supports a model in which BST-2 associates with SARS-CoV-2 structural proteins not only at the ERGIC and intracellular vesicles, but also at the plasma membrane, a potential site of virion release in certain cell types.

These results demonstrate that BST-2 physically associates with SARS-CoV-2 structural proteins at multiple subcellular locations. The spatial proximity between BST-2 and E, M, and N proteins is consistent with its known function as a tethering factor that impedes virion release by directly or indirectly anchoring nascent viral particles to cellular membranes. Together with our previous findings that BST-2 localizes to the ERGIC and restricts VLP release, these data support a model in which BST-2 acts at intracellular sites of viral assembly as well as at the cell surface to limit SARS-CoV-2 dissemination.

### SARS-CoV-2 ORF3a does not function as a BST-2 antagonist

Although prior studies have suggested that SARS-CoV-2 accessory protein ORF3a antagonizes the antiviral activity of BST-2 [31], our data do not support this function. To assess the potential role of ORF3a in BST-2 antagonism, we employed our established VLP release assay in BST-2-deficient HEK 293T cells. Co-expression of SARS-Cov-2 structural proteins (E, M, and N) with ORF3a led to enhanced VLP release compared to structural proteins alone. However, this enhancement occurred in the complete absence of BST-2 (data not shown), indicating that ORF3a’s ability to promote VLP release does not require or involve antagonism of BST-2 mediated restriction. These findings suggest ORF3a facilitates virus egress through mechanisms independent of BST-2.

To better understand the cellular context of ORF3a function, we examined its subcellular localization. HeLa cells were transfected with a plasmid encoding EGFP-tagged ORF3a and stained for ERGIC-53 to assess colocalization with the ERGIC. Although ORF3a-EGFP exhibited a perinuclear distribution, it showed minimal colocalization with ERGIC-53 (Pearson’s R = 0.29 ± 0.04), and ERGIC-53 morphology remained unchanged in the presence of ORF3a (Fig. 4a). These observations indicate that ORF3a does not accumulate at the ERGIC and likely functions within a distinct cellular compartment.

**Figure 4.**
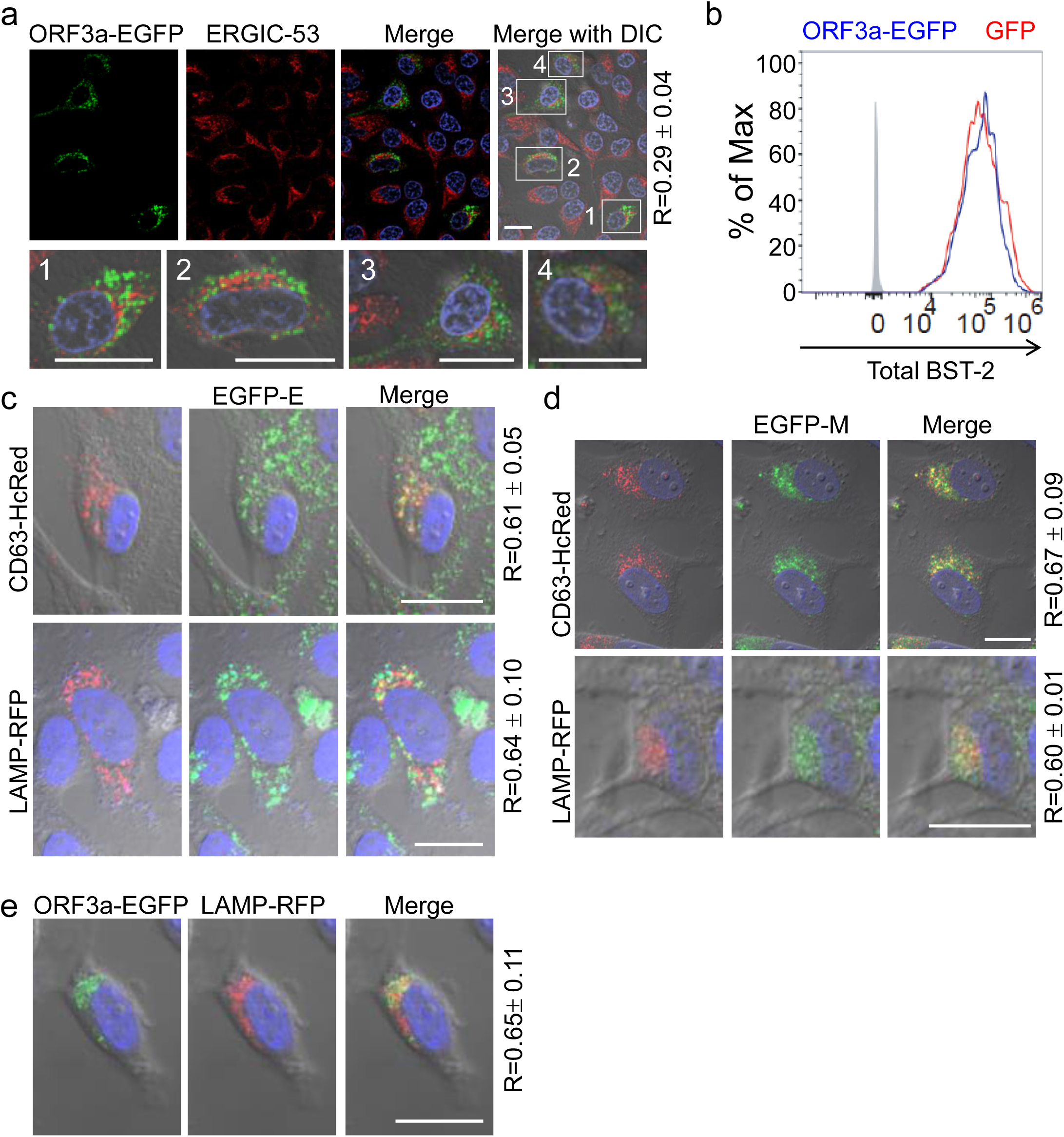
ORF3a does not localize to the ERGIC or alter BST-2 expression. **(a)** HeLa cells expressing ORF3a-EGFP were stained with anti-ERGIC-53. ORF3a-EGFP is shown in green, ERGIC-53 in red. Merged images and enlarged boxed regions are presented. **(b)** HeLa cells transfected with GFP or OFR3a-EGFP were stained for total BST-2 and analyzed by flow cytometry. Isotytpe control (gray), GFP-positive (red outline), and ORF3a-EGFP-positive (blue outline) populations are shown. **(c, d)** HeLa cells were co-transfected with EGFP-E (c) or EGFP-M (d) and either CD63-HcRed (top rows) or LAMP-RFP (bottom rows). **(e)** HeLa cells co-expressing ORF3a-EGFP and LAMP-RFP were imaged. ORF3a-EGFP is shown in green; LAMP-RFP in red. Yellow signal in merged images indicates colocalization. Scale bars, 20 μm. Pearson’s R values represent means ± SD from ≥ 3 independent experiments.

We next assessed whether ORF3a modulates BST-2 protein levels. Flow cytometric analysis of HeLa cells transfected with either ORF3a-EGFP or a control GFP construct revealed no significant difference in endogenous BST-2 levels (Fig. 4b). These data argue against a role for ORF3a in mediating the downregulation or degradation of BST-2, a mechanism commonly employed by viral antagonists such as HIV-1 Vpu [32].

Given these findings, we further explored alternative trafficking pathways in which ORF3a might localize and function. Confocal microscopy revealed strong colocalization of ORF3a-EGFP with LAMP-RFP, a lysosomal marker, with a Pearson’s R value of 0.65 ± 0.11 (Fig. 4e), indicating predominant localization within the late endosome-lysosome pathway. Similarly, EGFP-tagged E and M proteins also colocalized with late endosomal and lysosomal markers, including CD63-HcRed and LAMP-RFP, exhibited R values ranging from 0.60 to 0.67 (Fig. 4c, 4d). These results suggest that ORF3a and other SARS-CoV-2 structural proteins utilize late endosomal and lysosomal trafficking pathways, potentially facilitating non-canonical routes of viral egress.

Together, these findings demonstrate that ORF3a does not act as a BST-2 antagonist via degradation, downregulation, or sequestration. Instead, its role in promoting virus release likely involves the exploitation of endolysosomal trafficking pathways distinct from those regulated by BST-2. This function may be particularly important in cell types where viral egress is uncoupled from the conventional secretory pathway.

### ORF7a enhances SARS-CoV-2 release by promoting BST-2 downregulation

In contrast to ORF3a, the SARS-CoV-2 accessory protein ORF7a exhibited functional and localization properties consistent with BST-2 antagonism. To determine its subcellular localization, we expressed ORF7a in HeLa and HEK 293T cells and assessed colocalization with ERGIC-53. Confocal microscopy revealed a predominantly perinuclear distribution and strong colocalization with ERGIC-53 in both cell lines, with Pearson’s correlation coefficients of R = 0.73 ± 0.05 in HeLa cells and R = 0.73 ± 0.06 in HEK 293T cells (Fig. 5a). These findings are consistent with previous reports and support a role for ORF7a in modulating ERGIC-resident host restriction factors, such as SERINC5 [33] .

**Figure 5.**
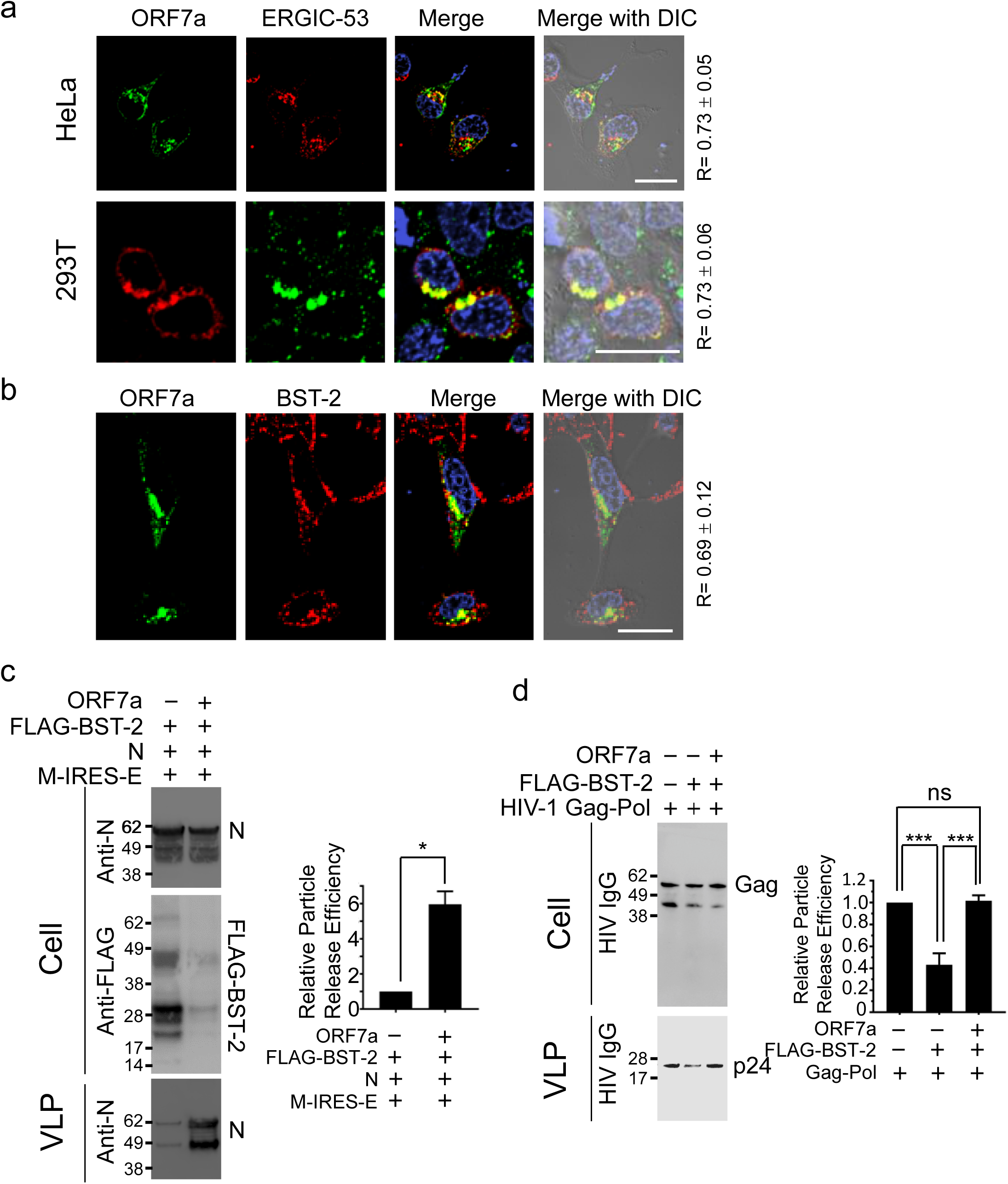
ORF7a functions as a BST-2 antagonist. **(a)** HeLa (top) and HEK 293T (bottom) cells expressing ORF7a were stained with anti-ORF7a and anti-ERGIC-53 antibodies. **(b)** HeLa cells expressing ORF7a were stained with anti-ORF7a and anti-BST-2. ORF7a is shown in green; ERGIC-53 or BST-2 in red. Merged DAPI and DAPI/DIC images are shown. **(c)** HEK 293T cells was co-transfected with M-IRES-E, N, FLAG-BST-2, and either empty vector or ORF7a. Cell lysates and VLPs were analyzed by western blot for N protein. Quantification of N in supernatants is shown below (means ± SD; n=3). *, *p* < 0.05. **(d)** HEK 293T cells were transfected with HIV-1 Gag-Pol alone, with FLAG-BST-2, or with both FLAG-BST-2 and ORF7a. Western blot analysis of VLPs and cell lysates were performed using HIV-1 IgG. Quantification of p24 in supernatants is shown below. ns, not significant; ***, *p* < 0.001. Scale bars, 20 μm. Pearson’s R values represent means ± SD from ≥ 3 experiments.

To assess the spatial relationship between ORF7a and BST-2, we performed colocalization analysis in HeLa cells using immunostaining for endogenous BST-2. ORF7a exhibited substantial overlap with BST-2 (R = 0.69 ± 0.12) (Fig. 5b), suggesting that these proteins reside in similar subcellular compartments and may physically interact or influence one another at the ERGIC or downstream vesicular compartments.

Functional assays confirmed the antagonistic activity of ORF7a toward BST-2. In the SARS-CoV-2 VLP release assay, co-expression of FLAG-tagged BST-2 significantly inhibited VLP release, as expected. However, co-expression of ORF7a restored VLP production to levels more than 5-fold higher than those observed in BST-2-expressing cells alone (Fig. 5c), demonstrating that ORF7a effectively counteracts BST-2-mediated restriction. Western blot analysis of cell lysates revealed that ORF7a expression led to a marked reduction in total cellular BST-2 levels, consistent with degradation or downregulation mechanisms commonly employed by viral antagonists.

To determine whether ORF7a’s antagonistic effect was specific to SARS-CoV-2 or represented a broader activity, we extended our analysis to an unrelated viral system. HEK 293T cells were transfected with HIV-1 Gag-Pol expression vectors and FLAG-BST-2, with or without ORF7a. In this context, BST-2 expression significantly reduced HIV-1 VLP release; however, co-expression of ORF7a rescued particle production to levels comparable to the vector control (Fig. 5d). These findings demonstrate that SARS-CoV-2 ORF7a exerts broad antagonistic activity against BST-2, independent of viral context.

Taken together, our results show that ORF7a localizes to ERGIC membranes where it colocalizes with BST-2, promotes its degradation or downregulation, and rescues virus-like particle release in both SARS-CoV-2 and HIV-1 systems. These observations establish ORF7a as a bona fide BST-2 antagonist that facilitates viral egress by neutralizing host restriction at key sites of virus assembly and trafficking.

### BST-2 restriction and ORF7a counteraction are functional in Calu-3 lung epithelial cells

To determine whether the BST-2-mediated restriction of SARS-CoV-2 release and its antagonism by ORF7a are operative in a physiologically relevant system, we extended our analysis to Calu-3 cells, a human lung epithelial cell line that endogenously expresses BST-2 and supports productive SARS-CoV-2 infection. Using our established VLP release assay, co-expression of the SARS-CoV-2 structural proteins E, M, and N in Calu-3 cells resulted in robust VLP production, confirming that these structural proteins are sufficient to drive particle assembly and egress in this system. However, the co-expression of FLAG-tagged BST-2 significantly suppressed VLP release, indicating that BST-2 retains its antiviral tethering function in lung epithelial cells. Importantly, co-expression of ORF7a reversed the suppressive effect of BST-2 and restored VLP release to near-control levels (Fig. 6a). These findings demonstrate that the antagonistic interaction between BST-2 and ORF7a is functional in Calu-3 cells, highlighting the physiological relevance of this host-virus interaction in the human respiratory epithelium.

**Figure 6.**
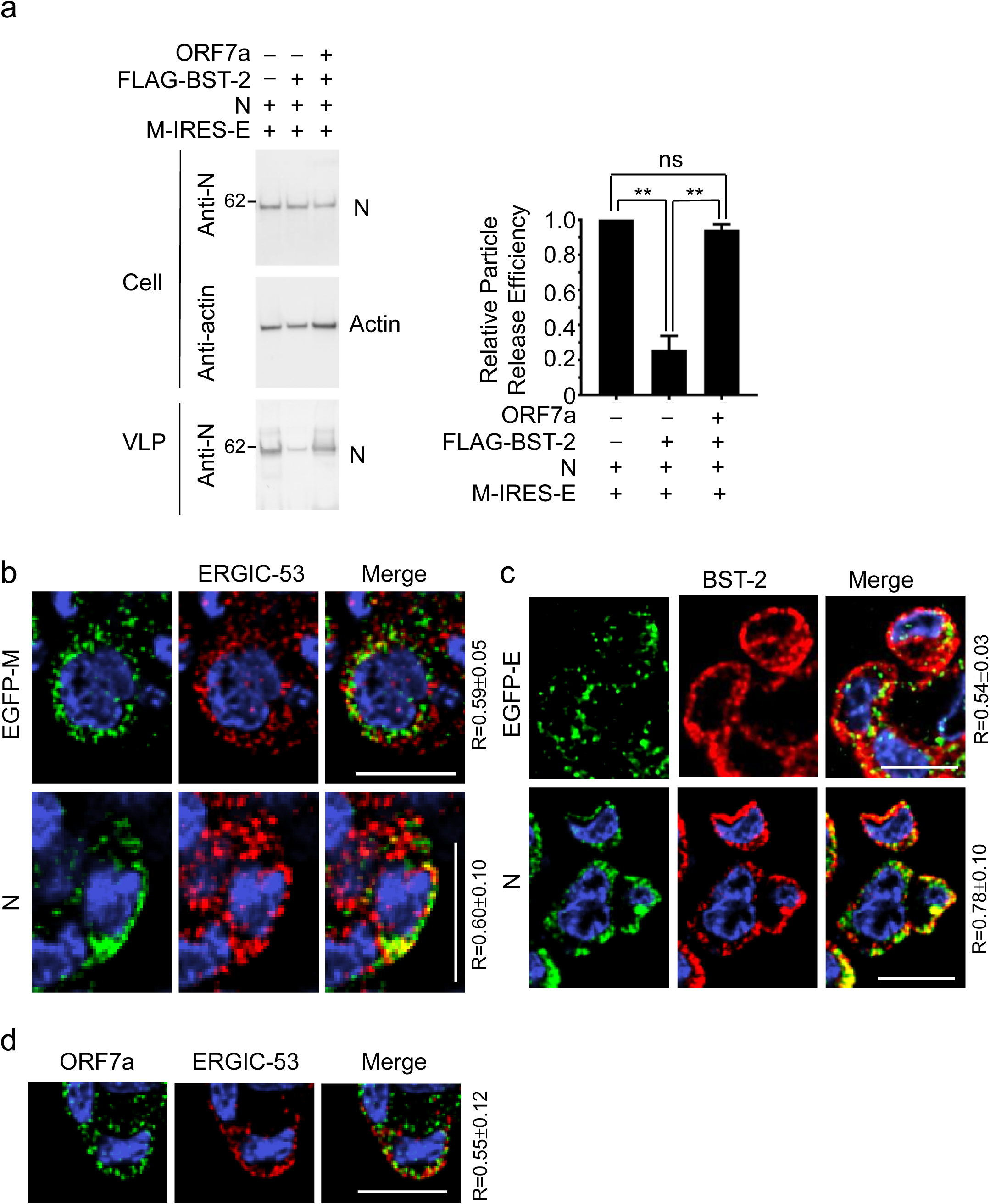
BST-2 restricts SARS-CoV-2 VLP release in Calu-3 cells and is counteracted by ORF7a. **(a)** Calu-3 cells were co-transfected with M-IRES-E and N alone, with BST-2, or with both BST-2 and ORF7a. Cell lysates and VLPs were analyzed by western blot for N protein. Quantification of N protein in supernatants is shown on the right (mean ± SD; n=3). ns, not significant; **, *p*<0.01. **(b)** Calu-3 cells transfected with EGFP-M (top) or N (bottom) were stained with anti-ERGIC-53 and anti-N antibodies. **(c)** Calu-3 cells transfected with EGFP-E (top) or N (bottom) were stained with anti-BST-2 and anti-N. **(d)** Calu-3 cells expressing ORF7a were stained with anti-ORF7a and anti-ERGIC-53. In (b-d), viral proteins are shown in green; cellular markers in red. Merged DAPI and DAPI/DIC images are shown in the right panels. Scale bars, 20 μm. Pearson’s R values represent means ± SD from ≥ 3 independent experiments with 20-30 cells per condition.

To further characterize the cellular context of ORF7a-mediated antagonism, we examined the subcellular localization of SARS-CoV-2 structural proteins and endogenous BST-2 in Calu-3 cells. Confocal microscopy revealed that EGFP-tagged M and N proteins colocalized with the ERGIC marker ERGIC-53, with Pearson’s correlation coefficients of R = 0.59 ± 0.05 and R = 0.60 ± 0.10, respectively (Fig. 6b). These values are consistent with those observed in HeLa and HEK 293T cells, supporting the conserved targeting of SARS-CoV-2 structural proteins to the ERGIC across multiple cell types.

We next assessed the spatial proximity of viral structural proteins to endogenous BST-2. Both EGFP-E and N exhibited measurable colocalization with BST-2, with Person’s R values of 0.54 ± 0.03 and 0.78 ± 0.10, respectively (Fig. 6c). The strong colocalization of N with BST-2 is particularly notable, as it suggests a close association at intracellular or plasma membrane sites, where BST-2 could restrict virion assembly or release at multiple stages of the viral life cycle.

Finally, we evaluated the subcellular localization of ORF7a in Calu-3 cells. ORF7a exhibited a partially perinuclear distribution and moderate colocalization with ERGIC-53 (R = 0.55 ± 0.12) (Fig. 6d). Although slightly lower than values observed in HeLa or HEK 293T cells, this level of colocalization supports the notion that ORF7a is appropriately positioned to antagonize BST-2 at or near the ERGIC in lung epithelial cells.

Collectively, these results demonstrate that both BST-2-mediated restriction and ORF7a-mediated counteraction are active in Calu-3 cells. They underscore the physiological relevance of this host-virus interaction and support a model in which SARS-CoV-2 evades host antiviral defenses at the ERGIC and plasma membrane to promote efficient viral egress in the respiratory epithelium.

## Discussion

In this study, we elucidate a mechanism by which the host restriction factor BST-2 inhibits the release of SARS-CoV-2 VLPs and demonstrate how the viral accessory protein ORF7a counteracts this restriction. Using a reconstituted VLP system composed of SARS-CoV-2 structural proteins M, E, and N, we found that co-expression of BST-2 significantly suppressed VLP release, consistent with its established role in physically tethering budding virions to cellular membranes and thereby preventing their dissemination. This antiviral effect was effectively reversed by co-expression of ORF7a, confirming ORF7a as a bona fide antagonist of BST-2. Confocal microscopy revealed BST-2 localization at both the plasma membrane and the ERGIC, indicating that BST-2 may impose antiviral barriers at multiple stages and cellular locales during virion egress. In contrast, ORF3a, previously implicated in BST-2 antagonism, continued to enhance VLP release in the absence of BST-2 under our experimental conditions, suggesting that its antagonistic effects are either context-dependent or mediated through alternative pathways distinct from direct BST-2 counteraction.

Our findings further reinforce the established model that the ERGIC serves as the principal site of coronavirus assembly and budding. Unlike many enveloped viruses that assemble at the plasma membrane, coronaviruses, including SARS-CoV-1 and SARS-CoV-2, utilize the ERGIC as a central hub for virion morphogenesis. Previous ultrastructural studies employing transmission electron microscopy and immunogold labeling have demonstrated the accumulation of budding virions within ERGIC-derived vesicles and the close association of viral proteins with ERGIC membranes [34–36]. Complementary biochemical fractionation and proteomic analyses support the enrichment of viral structural proteins and replicative intermediates within this compartment. Our immunofluorescence microscopy confirmed that all four SARS-CoV-2 structural proteins (E, M, N, and S) strongly colocalize with ERGIC-53, a canonical ERGIC marker, in both HeLa (Fig. 1a) and HEK 293T (Fig. 1b) cells. Quantitative analysis via Pearson’s correlation coefficients revealed particularly robust overlap for E and M proteins, consistent with their known roles as central drivers of membrane curvature and budding [37, 38]. The N protein exhibited partial colocalization, likely reflecting transient association mediated by interactions with M and genomic RNA packaging [39]. The S protein showed a more diffuse distribution, with distinct puncta overlapping ERGIC-53-positive regions, consistent with its synthesis in the ER and subsequent trafficking through the secretory pathway for virion incorporation. Moreover, strong colocalization among the structural proteins themselves (Fig. 1c, 1d) suggests that these proteins are not only targeted to the same compartment but also form pre-assembly complexes that coordinate envelopment and particle formation. These observations align with prior evidence identifying M as a scaffold for the recruitment of E, N, and S to assembly sites [37]. These interactions likely initiate in the ER and are stabilized within the ERGIC, which provides a unique membrane topology and protein environment conducive to coronavirus budding. Collectively, these results validate the use of ectopic expression systems to faithfully recapitulate coronavirus assembly and facilitate dissection of virus-host interactions.

BST-2 restricts a broad spectrum of enveloped viruses through a conserved mechanism involving dual membrane anchoring via its N-terminal transmembrane domain and C-terminal GPI anchor [40–42]. While its antiviral activity has been primarily characterized at the plasma membrane, emerging data indicate BST-2 also traffics through intracellular compartments, including endosomes and the TGN [4, 43]. Here, high-resolution confocal microscopy revealed that a significant fraction of endogenous BST-2 localizes to ERGIC-53-positive membranes in HeLa cells (Fig. 2b), situating BST-2 at the canonical site of coronavirus assembly. Notably, BST-2 robustly colocalized with SARS-CoV-2 structural proteins E and M, which are essential for viral envelope formation and budding from ERGIC membranes (Fig. 3a). These findings suggest that BST-2 is strategically positioned to restrict budding virions intracellularly, in addition to its plasma membrane function. Using a minimal VLP release assay lacking RNA and accessory proteins [44], we demonstrated that BST-2 co-expression significantly reduced particle release (Fig. 2a), confirming that BST-2’s antiviral effect occurs at the level of structural protein-mediated budding. This inhibition, independent of S protein, or genome replication, indicate direct interference with the budding process.

Mechanistically, these data support a model in which BST-2 retains fully or partially assembled virions at the ERGIC, either by physically tethering them to the limiting membrane of the budding compartment or by impeding vesicle-mediated transport toward the plasma membrane. This intracellular tethering extends BST-2’s antiviral repertoire beyond its classical role at the cell surface. Prior studies have reported BST-2-mediated intracellular restriction of hepatitis B virus budding into multivesicular bodies and inhibition of exosome secretion, both of which involve intracellular membrane compartments [45, 46]. However, the precise mechanisms underlying intracellular BST-2 activity remain to be elucidated, and whether these mechanisms mirror those at the plasma membrane is unclear. Given the ERGIC’s centrality in coronavirus assembly and egress through the secretory pathway, BST-2’s presence at this site likely poses a substantial barrier to viral release, particularly in interferon-stimulated cells. Our findings thus reveal a previously underappreciated facet of BST-2 antiviral function and broaden its scope to include the restriction of intracellularly budding viruses. This insight suggests that therapeutic modulation of BST-2 localization or stability could significantly impact viral release and innate immunity against coronaviruses.

Contrary to prior reports implicating ORF3a as a BST-2 antagonist [47], our data do not support such a role. Confocal imaging showed minimal colocalization of ORF3a with ERGIC-53 and no detectable effect on BST-2 expression or localization in HeLa cells (Fig. 4a, 4b). Importantly, ORF3a retained its ability to promote VLP release in BST-2-deficient HEK 293T cells (data not shown), indicating that its proviral function operates independently of BST-2 antagonism.

ORF3a predominantly localized to endolysosomal compartments, demonstrated by strong colocalization with LAMP1-positive late endosomes and lysosomes (Fig. 4e). Similar distribution patterns were observed for E and M proteins, indicating that SARS-CoV-2 exploits multiple intracellular trafficking pathways, including endosomal routes, alongside canonical ERGIC-based assembly and release (Fig. 4c, 4d). The inability of ORF3a to overcome BST-2 restriction suggests it facilitates viral egress via distinct mechanisms, consistent with emerging evidence implicating ORF3a in lysosomal exocytosis, autophagy modulation, and late endosome/lysosome trafficking [48–52]. These findings underscore the complexity of SARS-Cov-2 egress pathways, which may be differentially regulated by accessory proteins and vary between cell types.

Crucially, we demonstrate that ORF7a is a potent BST-2 antagonist capable of fully rescuing VLP release under BST-2 restriction (Fig. 5c). ORF7a colocalized with BST-2 and ERGIC-53 (Fig. 5a, 5b), localizing at or near the virion assembly site, and reduced BST-2 protein levels (Fig. 5c), consistent with targeted degradation. ORF7a also rescued HIV-1 VLP release under BST-2 restriction (Fig. 5d), indicating that it targets a conserved host antiviral mechanism shared across diverse viral families. These observations align with recent reports describing SARS-CoV-2 ORF7a-mediated BST-2 degradation [26]. The colocalization of ORF7a with BST-2 supports a model in which ORF7a sequesters or promotes degradation of BST-2 at the ERGIC, relieving its tethering effect and facilitating virion release. Although previous studies have documented direct ORF7a-BST-2 interactions [53–55], the precise degradation pathways utilized − whether lysosomal, proteasomal, and ER-associated degradation − remain to be elucidated.

Finally, we confirmed the physiological relevance of the BST-2/ORF7a axis in Calu-3 cells, a human lung epithelial model of SARS-CoV-2 infection. Here, structural proteins alone yielded robust VLP production, markedly inhibited by BST-2 and efficiently rescued by ORF7a (Fig. 6a). This demonstrates that BST-2-mediated restriction and ORF7a antagonism operate in a physiologically relevant context, particularly under interferon-induced antiviral states. By counteracting BST-2, ORF7a likely enhances viral egress and dissemination in airway epithelium, contributing to SARS-Cov-2 pathogenesis and transmission.

In summary, our study advances understanding of SARS-CoV-2 egress by identifying BST-2 as a multifaceted host restriction factor acting at intracellular and plasma membrane sites and establishing ORF7a as a critical viral antagonist. The distinct localization and BST-2-independent function of ORF3a highlight the existence of mechanistically diverse viral egress pathways. These insights reveal the complex interplay between viral accessory proteins and host antiviral defenses and offer potential targets for therapeutic strategies aimed at bolstering innate immunity to control coronavirus infection.

## Methods

### Cell lines and culture conditions

HEK 293T (CRL-3216), HeLa (CCL-2), and Calu-3 (HTB-55) cells were obtained from the American Type Culture Collection (ATCC). HEK 293T and HeLa cells were cultured in Dulbecco’s modified Eagle’s medium (DMEM; Thermo Fisher) supplemented with 10% fetal bovine serum (FBS), 100 U/ml penicillin, and 100 μg/ml streptomycin at 37°C in a humidified incubator with 5% CO2. Calu-3 cells were maintained in Eagle’s Minimum Essential Medium (EMEM; ATCC formulation) with 10% FBS under identical conditions.

### Plasmids

Expression plasmids EGFP-CoV M (Addgene #165124), EGFP-CoV E (Addgene #165123), and ORF3a-EGFP (Addgene #165121), encoding codon-optimized SARS-CoV-2 M, E, and ORF3a proteins with EGFP tags, respectively, were gifts from Bruce Antonny [56]. The plasmid pcDNA3.1-SARS-CoV-2 N (Addgene #158079), encoding the nucleocapsid (N) protein, was a gift from Jeremy Luban [57], while an alternative N construct (Addgene #153201) was obtained from Peter Klein [58]. The MAC-SARS-CoV-2 ORF7a plasmid (Addgene #158375), encoding a MAC-tagged ORF7a protein, was provided by Markku Varjosalo [59]. The CoV2-M-IRES-E plasmid (Addgene #177938), encoding codon-optimized M and E proteins separated by an IRES sequence, and the CoV2-N-WT-Hu1 plasmid (Addgene #177937), encoding codon-optimized N protein, were gifts from Jennifer Doudna[30]. The pcDNA3.1-SARS2-Spike plasmid (Addgene #145032), encoding a C9-tagged spike (S) protein, was a gift from Fang Li [60]. Additional SARS-CoV-2 S, M, and E expression plasmids (pcDNA3.1-hS, pcDNA3.1-hM, and pcDNA3.1-hE) were generously provided by Saveez Saffarian [61]. HA-tagged and FLAG-tagged BST-2 expression plasmids were provided by Vincent Piguet [62]. The HIV-1 Gag-Pol expression plasmid pGPCINS was obtained from Xiao-Fang Yu [63].

### Cell transfection

HEK 293T cells were transfected with either the polyethyleneimine (PEI, Sigma-Aldrich) or X-tremeGENE HP DNA Transfection Reagent (Sigma-Aldrich). HeLa cells were transfected using TransIT-HeLaMONSTER (Minus Bio), and Calu-3 cells were transfected with Lipofectamine 3000 (Invitrogen). In all cases, transfections were carried out following the manufacturers’ instructions.

### Primary antibodies

Mouse anti-Myc, anti-FLAG, and anti-ERGIC-53 antibodies were obtained from Sigma-Aldrich. Mouse anti-HA, and rabbit antibodies against SARS-CoV-2 nucleocapsid (N), spike glycoprotein (S), ORF3a, and ORF7a were obtained from Abcam. Mouse anti-BST-2 antibodies were purchased from Invitrogen. Rabbit anti-human BST-2 antibodies and HIV-1 IgG were obtained through the NIH HIV Reagent Program.

### Immunofluorescence microscopy

Immunofluorescence confocal microscopy was performed as described previously [64–66]. For single-staining experiments, ERGIC-53, BST-2, or HA-BST-2 was detected using mouse anti-ERGIC-53, anti-BST-2, or anti-HA antibodies, respectively, followed by goat anti-mouse Alexa Fluor 546-conjugated secondary antibodies (Life Technologies). For N protein single-staining, SARS-CoV-2 nucleocapsid (N) protein was detected using rabbit anti-N antibodies and goat anti-rabbit Alex Fluor 546-conjugated secondary antibodies (Life technologies). For double-staining experiments, ERGIC-53 was detected with mouse anti-ERGIC-53 antibodies and goat anti-mouse Alexa Fluor 546-conjugated secondary antibodies, while BST-2, HA-BST-2, N, or spike (S) protein was detected using rabbit anti-BST-2, anti-HA, anti-N or anti-S antibodies and goat anti-rabbit Alex Fluor 488-conjugated secondary antibodies. In experiments involving co-staining of BST-2 or HA-BST-2 with SARS-CoV-2 N, BST-2 or HA-BST-2 was detected with mouse anti-BST-2 or anti-HA antibodies and goat anti-mouse Alexa Fluor 546-conjugated secondary antibodies, while N protein was detected with rabbit anti-N antibodies and goat anti-rabbit Alex Fluor 488-conjugated secondary antibodies. For co-staining of BST-2 and SARS-CoV-2 ORF7a, BST-2 was detected using mouse anti-BST-2 antibodies and goat anti-mouse Alexa Fluor 546-conjugated secondary antibodies, while ORF7a was detected using rabbit anti-ORF7a antibodies and goat anti-rabbit Alex Fluor 488-conjugated secondary antibodies. Images were acquired using a Nikon A1R confocal microscope and analyzed using NIS-Elements AR software.

### VLP purification

HEK 293T or Calu-3 cells were co-transfected with SARS-CoV-2 M, E, and N expression plasmids, with or without co-expression of FLAG-BST-2 and /or ORF7a. For HIV-1 experiments, cells were co-transfected with pGPCINS, with or without FALG-BST-2 and/or ORF7a. Cell culture supernatants were collected 48 hours post-transfection, filtered through 0.45μm filters, and clarified by centrifugation at 3,000 rpm for 10 min at 4°C. VLPs were pelleted by ultracentrifugation through a 20% sucrose cushion at 28,000 × *g* for 2 hours at 4°C.

### Flow cytometry

HeLa cells transfected with either ORF3a-EGFP or GFP alone were harvested 48 hours post-transfection, fixed, and permeabilized. Cells were stained with mouse anti-BST-2 primary antibodies, followed by APC-conjugated goat anti-mouse IgG (H+L) secondary antibodies

(Invitrogen). Flow cytometric data were acquired using a BD FACScalibur flow cytometer (BD Biosciences) and analyzed using FlowJo software (BD).

## Acknowledgements

We thank Bruce Antonny. Jeremy Luban, Peter Klein, Markku Varjosalo, Jennifer Doudna, Fang Li, Saveez Saffarian, Vincent Piguet, Xiao-Fang Yu, Klaus Strebel, Amy Andrew, and Luiz Barbosa for providing reagents. We also thank Olga Korolkova and Qiujia Shao for technical assistance with confocal microscopy and flow cytometry.

## Funding

This research was supported in part by NIH grant R01AI1557764 (to X.D.), the Research Centers in Minority Institutions (RCMI) grant U54MD007586, and the Tennessee Center for AIDS Research (CFAR) grant P30AI110527.

### Author Contributions

X.D. conceived and designed the study. A.S. and X.D. performed the experiments. X.D. analyzed the data and wrote the manuscript. All authors reviewed and approved the final version of the manuscript.

